# High-depth whole genome sequencing of a large population-specific reference panel: Enhancing sensitivity, accuracy, and imputation

**DOI:** 10.1101/167924

**Authors:** Todd Lencz, Jin Yu, Cameron Palmer, Shai Carmi, Danny Ben-Avraham, Nir Barzilai, Susan Bressman, Ariel Darvasi, Judy H. Cho, Lorraine N. Clark, Zeynep H. Gümüş, Vijai Joseph, Robert Klein, Steven Lipkin, Kenneth Offit, Harry Ostrer, Laurie J. Ozelius, Inga Peter, Gil Atzmon, Itsik Pe’er

**Author notes:** **These two authors contributed equally** (Todd Lencz, Jin Yu).

## Abstract

**Background:** While increasingly large reference panels for genome-wide imputation have been recently made available, the degree to which imputation accuracy can be enhanced by population-specific reference panels remains an open question. In the present study, we sequenced at full-depth (≥30x) a moderately large (n=738) cohort of samples drawn from the Ashkenazi Jewish population across two platforms (Illumina X Ten and Complete Genomics, Inc.). We developed and refined a series of quality control steps to optimize sensitivity, specificity, and comprehensiveness of variant calls in the reference panel, and then tested the accuracy of imputation against target cohorts drawn from the same population.

**Results:** For samples sequenced on the Illumina X Ten platform, quality thresholds were identified that permitted highly accurate calling of single nucleotide variants across 94% of the genome. The Complete Genomics, Inc. platform was more conservative (fewer variants called) compared to the Illumina platform, but also demonstrated relatively greater numbers of false positives that needed to be filtered. Quality control procedures also permitted detection of novel genome reads that are not mapped to current reference or alternate assemblies. After stringent quality control, the population-specific reference panel produced more accurate and comprehensive imputation results relative to publicly available, large cosmopolitan reference panels. The population-specific reference panel also permitted enhanced filtering of clinically irrelevant variants from personal genomes.

**Conclusions:** Our primary results demonstrate enhanced accuracy of a population-specific imputation panel relative to cosmopolitan panels, especially in the range of infrequent (<5% non-reference allele frequency) and rare (<1% non-reference allele frequency) variants that may be most critical to further progress in mapping of complex phenotypes.

## Background

While genome-wide association studies (GWAS) have traditionally focused on the role of common genetic variation in common disease, it is increasingly acknowledged that rare genetic variation also significantly influences complex phenotypes [1]. The cost of sequencing to identify rare variation has declined dramatically in recent years; nevertheless, exome sequencing remains an order of magnitude higher in price compared to common SNP arrays, and high-depth whole-genome sequencing is even more expensive. Consequently, imputation has emerged as a popular approach that enables the examination of rare variants in the context of large-scale association studies in which subjects have been genotyped on conventional SNP arrays [2].

The accuracy and comprehensiveness of imputation depends on several factors, such as the size of the reference panel employed and the ancestry of its members relative to the target panel. Considerable effort has recently been devoted to generating increasingly large reference panels, usually involving cohorts sequenced at low-moderate (~4-10x) depth. Ancestry of these cohorts ranges from sampling across the global human population (e.g., 1000 Genomes Project Consortium [3]) to focusing on individuals from a particular population of origin such as the UK10K project [4] and the Genome of the Netherlands Consortium [5]. Attempts to characterize the trade-offs between breadth and depth of ancestry sampling have led to conflicting conclusions [6,7], although recent studies have consistently demonstrated enhanced accuracy of imputation for a given subpopulation when cosmopolitan reference panels such as 1000 Genomes are supplemented with data from population-specific reference panels [8,9]. Most recently, the Haplotype Reference Consortium [10] has assembled data from multiple worldwide studies involving a total of ~32,000 sequenced participants. It remains to be seen whether the utility of this resource can be further enhanced by population-specific sequencing.

In the present study, we tested whether imputation of rare variants could be improved by the addition of a moderately large (n=738), population-specific reference panel sequenced to full depth (≥30x). Specifically, we examined the Ashkenazi Jewish (AJ) population, which possesses unique characteristics, making it a compelling model for genetic investigation. While comprising ~10^7^ individuals today, the AJ population descends from a founding bottleneck that is very narrow, with effective population size likely less than a thousand chromosomes. This bottleneck is also very recent, having taken place only ~30 generations before present [11,12]. Thus, our reference panel conceivably samples each ancestral chromosome more than once at the average locus, allowing us to determine whether saturation of imputation can be achieved.

Development of enhanced referenced panels of sequenced individuals from a population also has relevance to the clinical setting, in which any personal genome includes many variants of unknown significance [13]. Such panels permit the identification of alleles that segregate in a population at appreciable frequencies, which otherwise might be erroneously interpreted as unique to a given patient with an unusual phenotype [14]. As with imputation, recent studies with increasingly large reference panels suggest that such distinctions remain problematic due to the extremely diffuse nature of human genetic diversity [15].

In developing our AJ reference panel, we sought to maximize the amount of information obtainable from each sample, by employing full-depth sequencing as well as a quality control (QC) pipeline that attempted to optimize the tradeoff between sensitivity (avoidance of false negative variant calls) and specificity (avoidance of false positive calls). To date, most QC pipelines for full-depth next-generation sequencing have sought to identify causal variants for unusual phenotypes, thus prioritizing the minimization of false positives, at times sacrificing regions of the genome that are more difficult to call. Similarly, recent consensus efforts to create “gold standard” reference calls tend to be limited to the most readily sequenced portions of the genome [16,17]. For example, it is widely acknowledged that low complexity regions (LCR), including homopolymers, short tandem repeats, and other repetitive elements comprise ~2% of the genome and are relatively inaccessible to accurate sequencing from short-read technologies [18].

Additionally, the 1000 Genomes Project divided the genomic space into three compartments (strict, pilot, and masked) based on observed read depth and mapping quality scores [3]. Only 77% of the genome was included in the ‘strict’ (highest-quality) compartment, defined as those regions in which local depth of coverage remained within 50% of the genome-wide average depth, and no more than 0.1% of reads have mapping quality of zero. By contrast, 4% of the genome demonstrated extremes of coverage (high and low) and contained many (>20%) low quality reads, and was therefore considered ‘masked.’ The remaining ‘pilot’ regions (19% of the genome) remained of questionable quality.

In the present manuscript, we describe steps that greatly decrease false positives and substantially increase the fraction of the genome that is confidently called. Additionally, because our reference panel was sequenced on two disparate platforms (Complete Genomics, Inc. (CGI) and Illumina Hiseq X Ten, see supplementary Figure 1), we describe an approach to reconcile data across platforms, extending previous work on this problem by including a larger number of replicate samples than previously described [19-22]. Finally, we identify new regions across the genome that are not currently well-mapped on either reference or alternate scaffolds, but which are routinely generated on the newest whole genome platforms, extending recent work on so-called blacklisted regions of the genome [23]. Thus, the present study is intended as a resource for the research and clinical genomics communities to enhance the interpretability of both large-scale genotype datasets and individual-level sequence data, while simultaneously providing a set of practical QC guidelines for end-users of the latest sequencing technologies.

**Figure 1.**
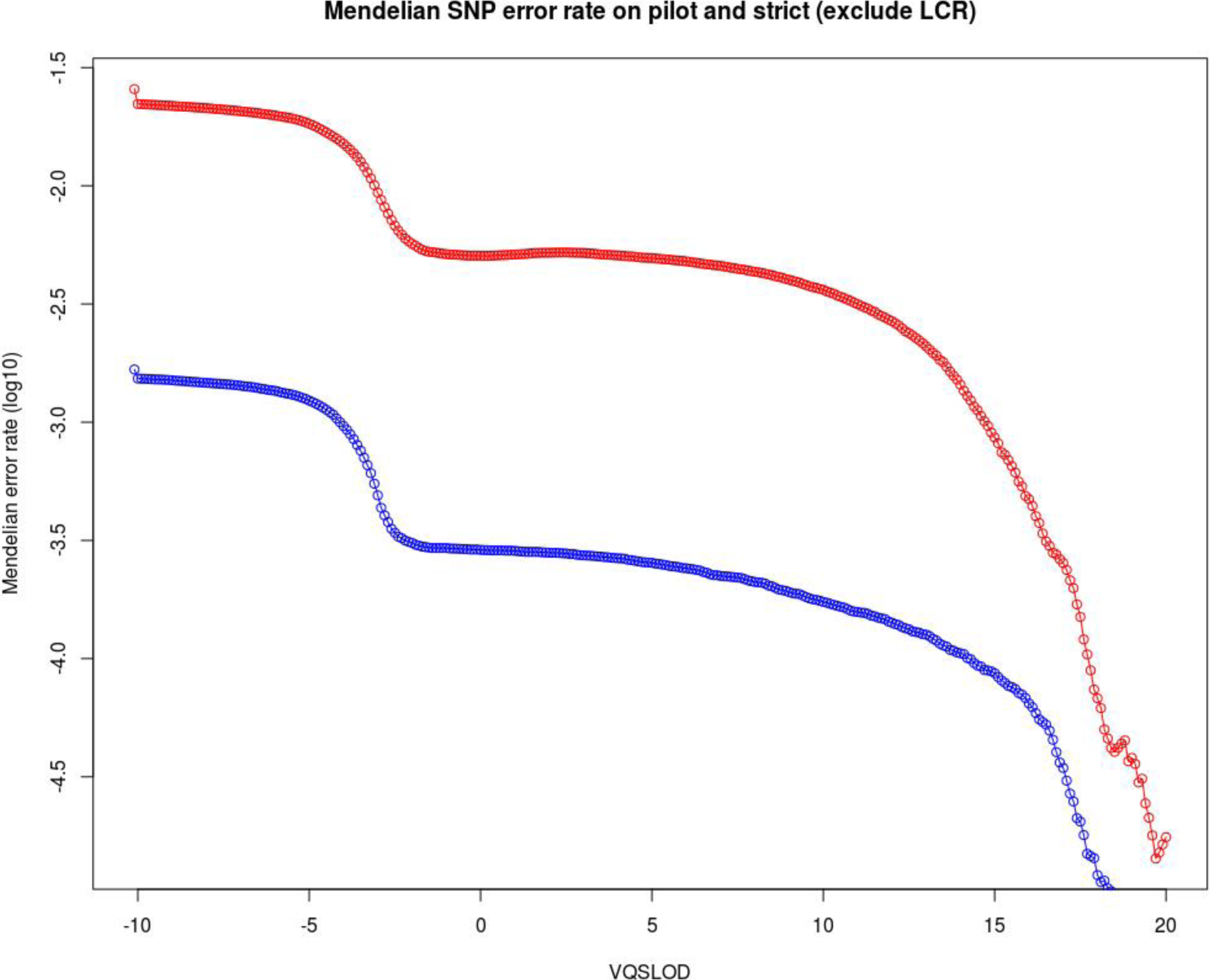
Observed Mendelian error rate (y-axis) as a function of reported quality (VQSLOD score, x-axis) for SNPs in non-LCR regions labeled as strict (blue) or pilot (red) calls. Both curves show rapid drop in error rate before a plateau starting at VQSLOD of −2.

## Results

### Enhancing call accuracy

After applying platform-specific filtering (see Methods), Mendelian errors in the Illumina-sequenced trio were reduced by nearly two orders of magnitude in both the pilot and strict non-LCR regions (Table 1a, cells marked in **bold**). Using default output from GATK (i.e., prior to our custom VQSLOD filtering), Mendelian error rates for single nucleotide variants (SNVs) were moderate (0.17%) in the strict, non-LCR region; however, across millions of variant sites, such a rate still implies several thousand false positives. Moreover, error rates were unacceptably high in all other compartments (Table 1a). Mendelian errors were reduced as VQSLOD filtering was increased, though in a non-linear fashion. As shown in Figure 1, a plateau was reached at VQSLOD=−2 for calls in the pilot region (red) and VQSLOD=−2.5 in the strict (blue) regions. Additionally, we applied a genotype quality (GQ) filter of 20, and call rate threshold of >90%, as variants below these thresholds were greatly over-represented in the masked and LCR regions.

The relationship between errors and VQSLOD was different for indels, such that no plateau was observed; true and false positives were intermingled across the quality spectrum (Supplementary Figure 2). Consequently, we utilized a VQSLOD=0 threshold, which resulted in a 1% Mendelian error rate for indels in the pilot and strict (non-LCR) compartments. When combined with GQ and call rate thresholds as above, filtering resulted in an acceptable Mendelian error rate (Table 1b). For both SNVs and indels, no usable relationships were observed between VQSLOD and error rates in LCR regions, or in the masked compartment so variants in these regions (~6% of the whole genome) were excluded from further analyses.

**Figure 2.**
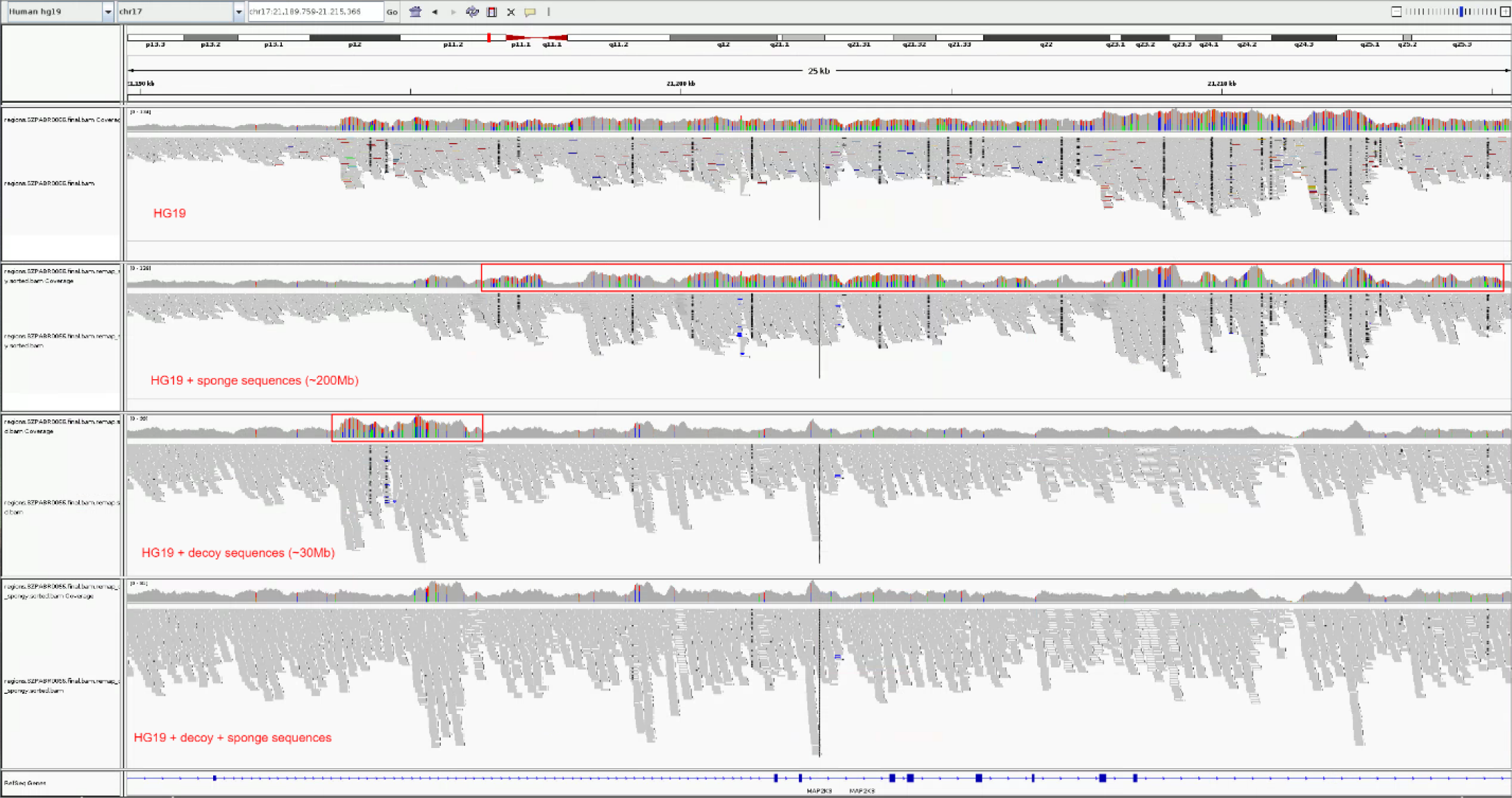
The utility of mapping to multiple reference sequences. Read-browser results of mapping for a 25kb segment along chromosome 17. Color identifies supposedly identified novel variants. The four panels from the top to bottom depict: 1) mapping against standard hg19 reference; 2) mapping against hg19 + “sponge” sequence of Miga et al. [23]; 3; mapping against hg19 + decoy sequences derived from the 1000 Genomes Project; 4; mapping against hg19 + sponge sequence + decoy sequence. Note that the two additional sets of reference sequences remove likely false-positive variants in different regions and combining them together could remove the most.

CGI samples (n=168) were called individually using a proprietary pipeline from Complete Genomics, Inc.; similar quality metrics were therefore not available. (It should be noted that n=36 of these CGI samples were held out for subsequent testing of the accuracy of the imputation reference panel.) False positives were identified by examination of nine samples run on both platforms, using Illumina data as the gold standard. Initial analysis of CGI data demonstrated that 7.99% of SNV calls were not observed in the Illumina calls for the same individuals. However, more than half (51.2%) of these false positives were observed to be singletons in the full CGI dataset. Consequently, we applied the following filters to the CGI data: 1) remove singletons; 2) remove masked and LCR regions; 3) remove variants called in <90% of CGI samples. Of the resulting SNVs calls, 97.2% were validated in the filtered Illumina data. Similar results were obtained in the CGI indel data; after applying the same filtering pipeline, however, only 91.5% were validated in the filtered Illumina data. Consequently, any indels which were only observed on the CGI platform (and never in the Illumina dataset) were filtered.

### Cross-Platform Merging

In total, we observed 17.6M variants in the filtered Illumina dataset, and around a half (~8.8M) of these in the smaller CGI dataset. Among the SNVs called in both Illumina and CGI data, virtually all (99.99%) had consistent allele frequencies across platform. Only 918 SNPs had allele frequency differences > 0.2 (Supplementary Figure 3), which would correspond to >6 standard deviations for a randomly sampled common variant. Outlying frequencies of Illumina variants that were inconsistent with their CGI frequencies were often small-integer fractions, suggestive of copy-number artifacts and motivating cross platform filtering (see Methods, and Supplementary Table 1, first three rows).

**Figure 3.**
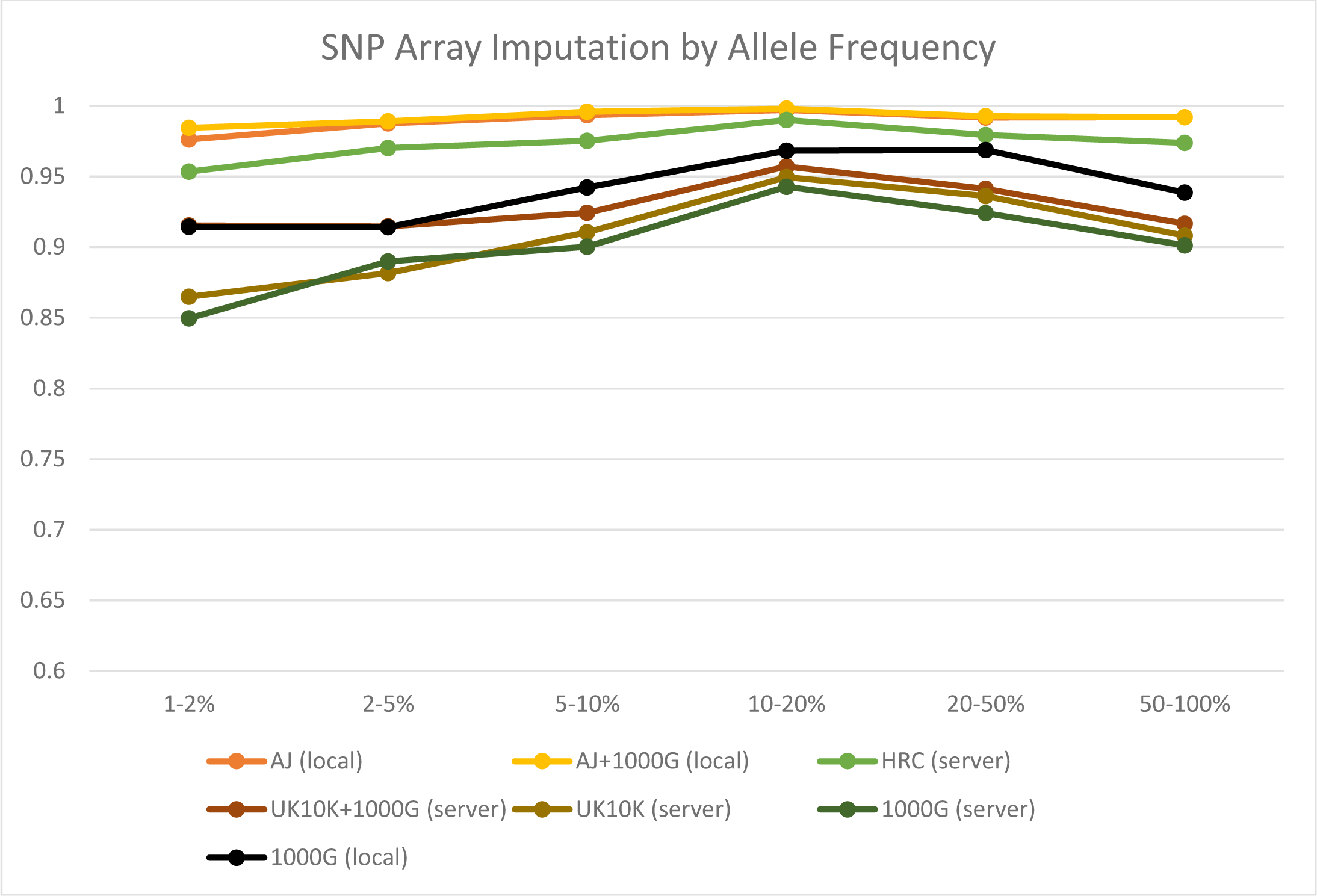
Imputation accuracy (aggregated R^2^; y-axis) across non-reference allele frequencies (x-axis) at held-out SNP-array sites in a genotyped panel of 2195 AJ individuals. Imputation quality is high for common alleles, but rarer ones were imputed markedly better using AJ references samples (AJ only – green, AJ combined with the 1000 genomes, navy blue).

**Table 1.** Mendelian Errors by Compartment and Filtering

| *Repeat Region* | LCR |  |  | Non-LCR |  |  |
| --- | --- | --- | --- | --- | --- | --- |
| *Compartment* | Masked | Pilot | Strict | Masked | Pilot | Strict |
| *Unfiltered* | 14.66% | 14.72% | 2.83% | 7.89% | 2.57% | 0.17% |
| *Post-Filter* | N/A |  |  |  | **0.0415%** | **0.0049%** |

| *Repeat Region* | LCR |  |  | Non-LCR |  |  |
| --- | --- | --- | --- | --- | --- | --- |
| *Compartment* | Masked | Pilot | Strict | Masked | Pilot | Strict |
| *Unfiltered* | 10.80% | 10.75% | 14.77% | 10.23% | 3.54% | 1.51% |
| *Post-Filter* | N/A |  |  |  | **0.1756%** | **0.0829%** |

Notably, we found that high-frequency Illumina-only SNVs that were not observed in the 1000 Genomes database tended to cluster within specific chromosomal intervals, often (but not always) near the telomeres and centromeres (Supplementary Figure 4). One such region is displayed in Figure 2; in the top panel, a lengthy stretch of uncatalogued variants is observed in a 25kb segment within the gene *MAP2K3*, near the chromosome 17 centromere. As shown in the lower panels, mapping against the alternate “sponge sequence” developed by Miga and colleagues [23] and the “decoy” sequence proposed by the 1000 Genomes Consortium [3] re-assigns these reads, such that these variants are no longer mapped within this gene sequence. However, not all such regions were completely cleared using either of these published alternative scaffolds, nor using the hg38 alignment. We identified 90 regions that harbor high-frequency, uncatalogued runs of variants as listed in Supplementary Table 2. Copy number phenomena are a plausible source of these anomalies, given that these regions are marked by individuals carrying derived alleles, called heterozygous at multiple neighboring positions. However, the boundaries of these regions do not precisely track boundaries of known copy number variants in DGV. For purposes of developing the imputation reference, SNPs in these regions were filtered using a loose Hardy-Weinberg threshold (p<10^-10^), yielding a final set of 17.5M SNVs in our combined reference dataset (total N=738).

**Figure 4.**
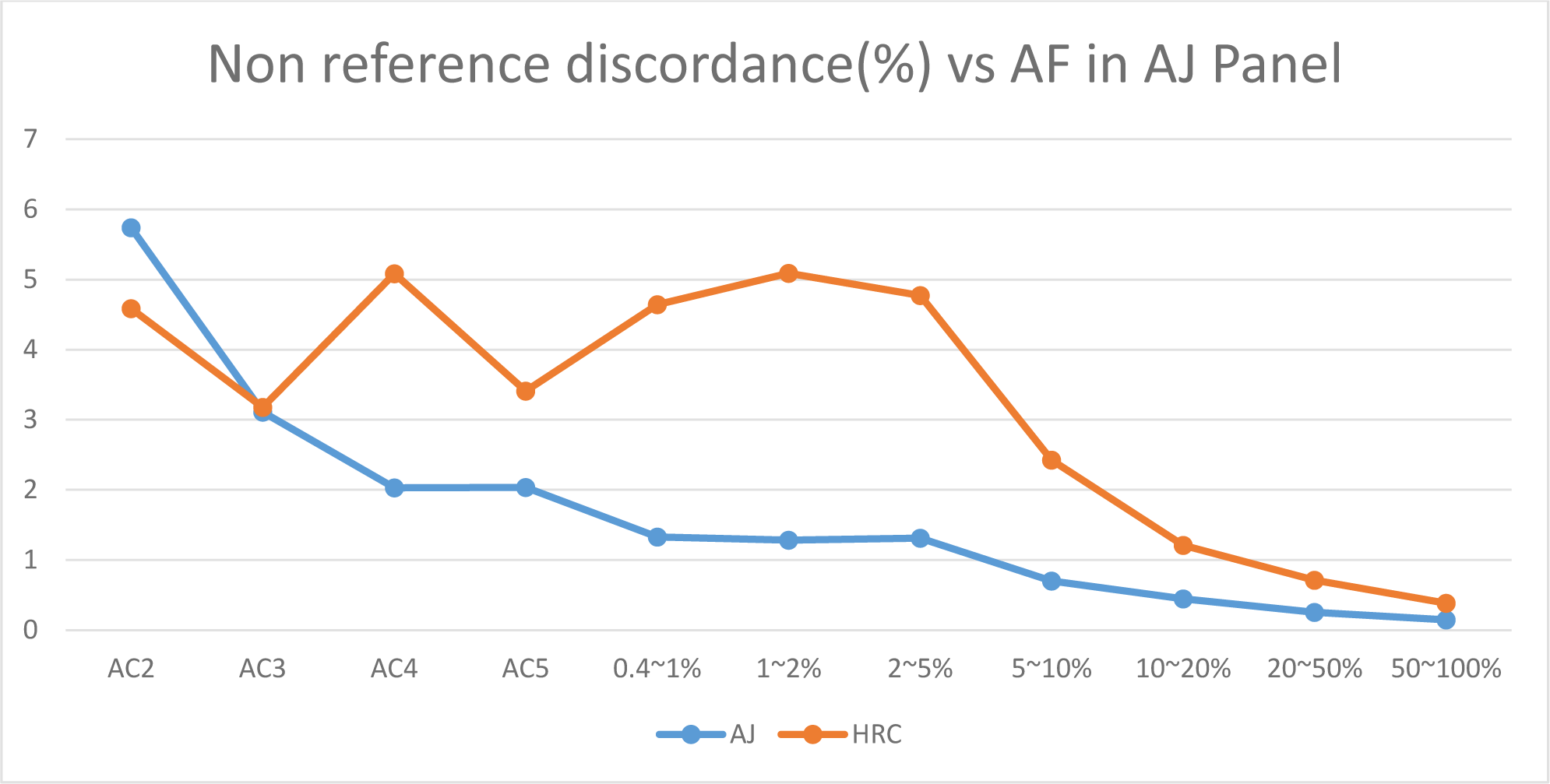
Non-reference genotype discordance (y-axis) across non-reference allele frequencies (x-axis) at overlapping variant sites between AJ reference panel and HRC reference panel in a sample of 36 AJ individuals sequenced on the CGI platform. Note that the non-reference allele counts/frequencies were calculated in the AJ panel.

**Table 2.** Imputation accuracy

| Panel | Pipeline | RR | RA | AA | NDR |
| --- | --- | --- | --- | --- | --- |
| **1000 Genomes** | Local | 0.44 | 1.94 | 1.43 | 2.69 |
| **1000 Genomes** | Server | 0.69 | 3.16 | 2.17 | 4.29 |
| **UK10K** | Server | 0.60 | 3.29 | 2.23 | 4.14 |
| **1000G + UK10K** | Server | 0.48 | 2.63 | 2.02 | 3.40 |
| **HRC** | Server | 0.19 | 0.85 | 0.58 | 1.15 |
| **AJ sequencing** | Local | 0.08 | 0.26 | 0.30 | 0.47 |
| **AJ + 1000G** | Local | **0.06** | **0.34** | **0.17** | **0.40** |

### Imputation Performance

We compared the contribution of our newly constructed population-specific reference panel to the accuracy of imputing common and infrequent alleles in our sample of n=2195 Ashkenazi subjects genotyped on the Omni-Quad chip array (see Methods). As shown in Table 2, we observe discordance rates to be nearly an order of magnitude smaller when using the Ashkenazi reference sequences as compared to the cosmopolitan 1000 Genomes panel and the European-specific UK10K panel. Performance of the AJ panel is ~2-fold better than the cosmopolitan HRC panel, despite a difference in sample sizes that is two orders of magnitude. Improved performance of the AJ-specific panel is especially meaningful at more rare alleles (Figure 3), where imputation using a cosmopolitan panel is often subpar. We observed slight improvement when we combined the AJ-specific panel with the 1000 Genomes panel.

We also evaluated all overlapping sites between AJ panel and HRC panel on chromosome 20 using a sample of 36 AJ individuals sequenced on the CGI platform as the gold standard (Methods). Overall, the non-reference genotype discordance rates of AJ panel and HRC panel are 0.32% and 0.97% respectively, which corresponds to a 3-fold improvement in AJ panel over HRC panel (Table 3, top two rows). This evaluation includes a much larger number of variants and a wider spectrum of allele frequency compared to the SNP array analysis. The non-reference genotype discordance rates were then plotted versus allele frequency bins (Figure 4). The AJ panel outperformed the HRC panel across all allele frequency bins above 0.145% (i.e., allele count=2 in the AJ reference panel, Figure 4), despite the HRC panel having a much larger sample size and minimal allele counts of 5 for all the overlapping variants.

**Table 3.** Imputation accuracy for CGI hold-out samples

|  | NRD (%) | RR(%) | RA->RR | AA->RR | RA(%) | RR->RA | AA->RA | AA(%) | RR->AA | RA->AA |
| --- | --- | --- | --- | --- | --- | --- | --- | --- | --- | --- |
| **AJ overlap** | 0.32 | 0.06 | 0.06 | 0.00 | 0.23 | 0.16 | 0.07 | 0.13 | 0.00 | 0.13 |
| **HRC overlap** | 0.97 | 0.22 | 0.21 | 0.00 | 0.62 | 0.45 | 0.16 | 0.36 | 0.01 | 0.35 |
| **HRC only** | 2.76 | 0.21 | 0.21 | 0.00 | 3.97 | 3.67 | 0.30 | 0.24 | 0.02 | 0.22 |
| **AJ only (CGI and/or Illumina)** | 13.66 | 0.13 | 0.13 | 0.00 | 17.63 | 17.48 | 0.15 | 0.78 | 0.48 | 0.30 |
| **AJ only - CGI only** | 96.95 | 0.36 | 0.35 | 0.01 | 96.67 | 96.64 | 0.04 | 98.75 | 97.18 | 1.41 |
| **AJ only - Illumina only** | 1.47 | 0.05 | 0.05 | 0.00 | 1.48 | 1.30 | 0.18 | 0.30 | 0.01 | 0.30 |

Given the biological importance of rare variants, we also evaluated the imputation accuracy of the rarest variants in our constructed AJ panel, which are also most likely to be private in AJ population and not called in HRC panel (Methods). For variants observed at 0.36% frequency (allele count = 5) in the AJ reference cohort, many individual samples show false positive and negative rates of 0% (Supplementary Figure 5, top panel). Notably, a larger proportion of imputed variants received a no-call in the CGI sequence data, due to its known conservative bias [19,21]. Notably, the overwhelming majority of false positive imputation calls were drawn from variants that were observed exclusively in the CGI samples of the reference panel, and were not present in the Illumina-sequenced samples of the AJ reference panel or in the HRC panel (Table 3, bottom three rows). Consequently, as a final step in construction and cleaning of the AJ reference panel, we eliminated all such variants exclusive to the CGI samples (Supplementary Table 1, bottom row).

**Figure 5.**
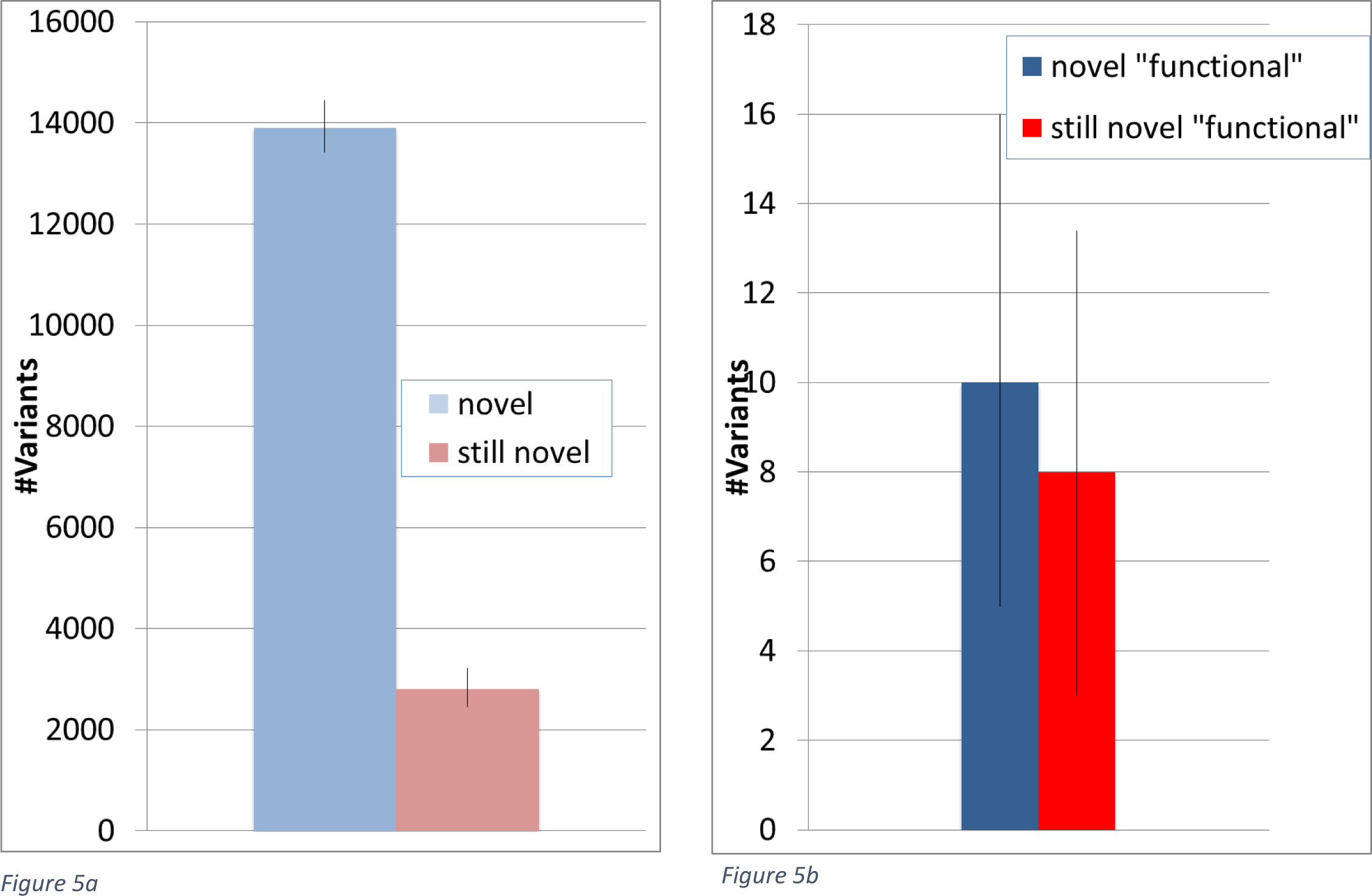
Variants in an out-of-sample AJ personal genome deemed novel before (blue) and after (red) inclusion of the dataset reported in this study. Data is presented for the median held-out sample (errorbars: 90%-CI), for all variants genome-wide (panel a) or only functional (exonic/splicing) ones (panel b).

### Population-specific variant discovery

Finally, we sought to evaluate the potential utility of the population-specific reference panel as a catalogue of normal variation in the population, applicable to the clinical interpretation of personal genomes (see Methods). Filtering against the remaining reference panel removed nearly 80% of novel variants across the entire genome (Figure 5a). When considering only novel functional (coding or splicing) variants, the population specific panel improves filtering significantly (p<1.1*10^−10^), but less dramatically, reducing the median count of such variants from 10 to 8 (Figure 5b).

## Discussion

Our primary results (Tables 1 and 3; Figures 3 and 4) demonstrate enhanced accuracy of a population-specific imputation panel relative to cosmopolitan panels, especially in the range of infrequent (<5% non-reference allele frequency) and rare (<1%) variants that may be most critical to further progress in mapping complex phenotypes [24,25]. These results extend prior studies that have shown the superiority of combining population-specific with cosmopolitan panels [8,9], by demonstrating: 1) a moderately-sized population-specific reference sample sequenced to full-depth provides better performance than the newly-released Haplotype Reference Consortium panel [10]; and 2) addition of a cosmopolitan panel (e.g., 1000 Genomes) to such a population-specific panel provides only marginal improvement in performance, consistent with recent findings in an outbred population cohort [26].

Moreover, for clinical purposes of interpreting a personal genome, our population-specific panel significantly enhanced filtering of variants unlikely to be related to disease (Figure 5). However, there is still considerable room for additional filtering in the coding regions (Figure 5b). As the exome is under greater pressure from negative selection compared to the rest of the genome, it is likely that many exonic variants are of relatively recent origin (i.e., post-dating the bottleneck in AJ history). Given the rapidly expanding nature of the human population in recent centuries (including the post-bottleneck AJ population), exceptionally large samples will be needed to achieve asymptotic representation of background variation [27,28]. In the applied setting, sequencing of parents may be the most efficient strategy for filtering and variant interpretation [29], supplemented by large-scale sequencing resources from the general population [30] and moderate-scale population-specific resources such as those described here.

While providing benefits such as those described above, short-read sequencing technologies continue to have technical challenges and limitations that we have sought to address in the present study. While many studies [31,32] have compared accuracy of different alignment and mapping protocols for raw short-read data, such comparisons are computationally expensive and may be impractical for application to large population cohorts and unavailable in the clinical setting. Many end-users of sequencing data receive batched calls from sequencing centers using standard workflows using the Genome Analysis Toolkit (GATK [36]). In this context, the present report provides practical guidelines to filtering for such users (both research and clinical). Our approach yields high accuracy not only for the most stably called regions, but also the so-called “pilot” region, extending high accuracy variant calling to 94% of the genome. This compares favorably to recent approaches such as the Genome in a Bottle [16,17] and ReliableGenome [33], which primarily focus on the optimizing accuracy of the ~70% of the genome that is least susceptible to technical artifact in short-read data.

The potential clinical importance of expanding the range of the genome that can be called reliably is illustrated in Supplementary Figure 6. In this figure, we parse the genes designated by the ACMG as harboring clinically actionable variants [34] as a function of proportion of variants observed in each calling compartment (strict/pilot/masked/LCR) as designated by the 1000 Genomes Project. Each of these genes contains segments that fall in the pilot compartment (Supplementary Figure 6a), and several genes (such as *PMS2, SDHC*, and *SDHD*) contain up to 50% of exonic bases designated as pilot. Thus, a clinical readout that is unable to accurately capture these bases would be relatively incomplete.

Several limitations in the short read data were difficult to overcome. For example, while we were able to produce reasonable error rates for indels even within the pilot compartment, these required more careful filtering, and do not have a readily identifiable optimum for balancing false positives vs. false negatives (Supplementary Figure 2). Additionally, we observed that the CGI platform suffers from two significant limitations: it is generally overly conservative (fewer total calls, more non-calls) compared to Illumina, but at the same time it is susceptible to a relatively large number of platform-specific false positives. While this platform is no longer active, legacy datasets should be treated with caution for novel variants, although known variants are conservatively called. Finally, we have identified regions of the genome that are not well mapped in current reference or alternate assemblies (Figure 2; Supplementary Figure 4). Further work is needed to properly characterize the (likely) structural variations that underlie these anomalous segments [35].

## Methods

### Cohort description

Samples sequenced using CGI (n=168) included control samples (n=128) described previously[11] as well as schizophrenia cases (n=40). Illumina samples (n=574) included samples described previously from multiple case-control cohorts described in Supplementary Table 3. By design, these samples included some overlapping subsets CGI controls (n=5) and CGI schizophrenia cases (n=4). All samples had been self-reported to be Ashkenazi Jewish, and also previously genotyped by SNP arrays, thus verified as Ashkenazi Jewish by principal components analysis of SNP data.

### Sequencing and analysis pipeline

For samples sequenced on the Illumina platform, genomic DNA was isolated from whole blood and was quantified using PicoGreen on a Spectramax fluorometer (Molecular Devices) or Qubit (Life Technologies), and integrity assessed using the Fragment Analyzer (Advanced Analytical). A separate aliquot was removed and used for SNP array genotyping using the HumanExome-12 v1.2-A (Illumina) chip. Sequencing libraries were prepared using the Illumina TruSeq Nano DNA kit, with 100ng input gDNA following manufacturer recommendations. Briefly, DNA was first sheared on a Covaris sonicator, followed by end-repair of the fragmented molecules and bead based size selection. Size selected molecules were then A-tailed followed by ligation of sequencing adaptors. Libraries were evaluated using a BioAnalyzer (Agilent), and quantified by qPCR (Kappa) and PicoGreen.

Libraries were sequenced on the Illumina HiSeq X ten with v1 chemistry. Sequencing libraries were pooled in equimolar amounts (8 samples / pool), and 2.5nM or 3nM pooled library was loaded onto each lane of the patterned flow cell, and clustered on a cBot (Illumina). Each pool of libraries was sequenced on 8 lanes of a flow cell. The HiSeq X generates ~375-400M pass filter 2x150bp per flow cell lanes. For samples that did not meet 30x mean genome coverage post alignment, additional aliquots of the sequencing libraries were pooled in proportion to the amount of additional reads needed, and re-sequenced in one or more flow cell lanes.

Upon completion of sequencing run, bcl files were demultiplexed and quality of sequencing data reviewed using SAV software (Illumina) and FastQC (http://www.bioinformatics.babraham.ac.uk/projects/fastqc/) for deviations from expected values with respect to total number of reads, percent reads demultiplexed (>95%), percent clusters pass filter (>55%), base quality by lane and cycle, percent bases >Q30 for read 1 and read 2 (>75%), GC content, and percent N-content. FastQ files were aligned to GRCh37 using the Burrows-Wheeler Aligner (BWA-MEM v0.78) [37] and processed using the best-practices pipeline that includes marking of duplicate reads by the use of Picard tools (v1.83, http://picard.sourceforge.net), realignment around indels, and base recalibration via Genome Analysis Toolkit (GATK v3.2.2) [36].

Single nucleotide variants were called using GATK HaplotypeCaller, generating a single sample GVCF file. Batches of samples were jointly genotyped GATK GenotypeGVCFs to generate a multi-sample VCF. Variant Quality Score Recalibration (VQSR) was performed on the multi-sample VCF, and variants were annotated using VCFtools [38] and in-house software.

Sequencing procedures for Complete Genomics, Inc. have been described in our prior publication reporting on a subset of these samples [11] using procedures described previously [39,40]. The average raw sequencing depth was 56x. The first 58 genomes were called using CG pipeline 2.0.2.26. All other genomes were called using pipeline 2.0.4.14. Both pipelines mapped variants to reference genome version hg19.

### Platform-specific Filtering

The sensitivity-specificity tradeoff in detecting variants is a key step in large scale sequencing efforts, often requiring dataset-specific adjustments. To examine false positive rates, Mendelian errors for a trio included in the Illumina-sequenced batch were examined as a function of VQSLOD score and genomic compartment. Since the *de novo* mutation rate is ~1.6x10^-8^ per base pair per generation [41,42], this was considered negligible relative to the potential error rate.

### Cross-platform filtering

We filtered out ~5K SNVs with observed allele frequencies of >0.2 in the CGI dataset that were not observed at all in the larger Illumina dataset, as well as ~39K Illumina SNVs with allele frequency >0.2 that were not observed in either the CGI dataset or the 1000 Genomes database. We also filtered 916 SNVs which were called in both CGI and Illumina but had an allele frequency difference larger than 0.2, and an additional 483K SNPs not called in either Illumina batch or the HRC but contributes to majority of the imputation errors (Supplementary Table 1). We also filtered SNP and INDELs pass Hardy-Weinberg threshold (p<10^-10^) in at least one platform as the site filtering method used in constructing the HRC panel [10]. The multi-allelic variants are also filtered after the merging Illumina and CGI call sets for constructing the AJ reference imputation panel.

### Imputation Evaluation

We considered the accuracy of imputation with our population-specific reference panel by comparing it to large, cosmopolitan reference panels in three ways. First, we examined imputation of common and infrequent variants(>5% and >1% non-reference allele frequencies, respectively) utilizing an independent test cohort of AJ subjects (n=2195, after removing 349 subjects overlapping with the sequencing cohort) with high quality genotypes available at ~1M SNPs assayed using the Illumina Omni-Quad platform, as described previously [43]. A random subset of variants in the array data that were spread across the frequency spectrum were masked and held out to test concordance of imputed vs. masked genotypes. (Of course, it should be noted that some small percentage of genotyping errors exist in the array data.) Imputation was performed either locally, using SHAPEIT2 and Impute2, or using the HRC imputation server at Sanger (using SHAPEIT2 for pre-phasing and PBWT for imputation). Then genotype discordance (Table 2) and aggregated R^2 (Figure 2) were calculated respectively using the SNP array genotypes as the gold standard on chromosome 20.

Second, we evaluated the imputation accuracy of all the sites available on both the constructed AJ panel and the HRC panel on chromosome 20, which represents a wider allele frequency spectrum than the SNP array comparison. The genotypes of 36 AJ individuals sequenced by CGI (hold-out sample) were used as the gold standard to calculate the non-reference genotype discordance vs the allele frequency (Table3; Figure 4). Notably, we also evaluated the imputed genotypes on variants private to the AJ panel (i.e., not called in the HRC panel) and found the overwhelming majority of false positive imputation errors were drawn from variants that were observed exclusively in the CGI samples but not supported by the larger set of Illumina-sequenced samples of the AJ reference panel (Table 3). Consequently, as a final step in construction and cleaning of the AJ reference panel, we eliminated all such variants exclusive to the CGI samples (Supplementary Table 1, bottom row).

Last, we evaluated the imputation accuracy of the rarest variants in AJ panel, most of which are private in AJ population. To control the sequencing error at the minimal level, we evaluated all the variants with a non-reference allele count from 2 to 5 in Illumina sequencing samples (N=574) on chromosome 20. The genotypes of 36 AJ individuals sequenced in CGI were used as the gold standard. The absolute numbers of each comparison group are plotted for each of the 36 AJ individual. As expected, all of the imputed genotypes of the rarest variants are Ref/Ref and Ref/Alt. The false positive errors are represented by “impute_RA_RR” group which mean the true Ref/Ref genotypes were imputed to Ref/Alt, while the false negatives errors are represented by “impute_RR_RA” and “impute_RR_AA” groups. (Supplementary Figure 5)

### Evaluation of Variant Filtering in Personal Genomes

We employed a leave-one-out approach, where we examined novel (i.e., not in dbSNP147) variants in each individual sample in relation to the remaining samples in the reference panel. In a clinical context, such variants may be labeled variants of uncertain significance (VUS).

## Acknowledgements

We acknowledge financial support from the Human Frontier Science Program (SC); NIH research grants AG042188 (GA), DK62429, DK062422, DK092235 (JHC), NS050487, NS060113 (LNC), AG021654, AG027734 (NB), MH089964, MH095458, MH084098 (TL), and CA121852 (computational infrastructure, IPe’er); NSF research grants 08929882 and 0845677 (IPe’er); Rachel and Lewis Rudin Foundation (HE); North Shore - LIJ Health System Foundation (TL); Brain & Behavior Foundation (TL); US-Israel Binational Science Foundation (TL, AD); LUNGevity Foundation (ZHG); New York Crohn’s Disease Foundation (IPeter); Edwin and Caroline Levy and Joseph and Carol Reich (SB); the Parkinson’s Disease Foundation (LNC); the Sharon Levine Corzine Cancer Research Fund (KO); and the Andrew Sabin Family Research Fund (KO).

## Author Contributions

TL and IP led the analysis, and led the writing of the manuscript. JY, CP, and SC conducted the primary analyses. TL led the funding of the study. TL, AD, GA, DB, NB, and LNC provided samples and conducted lab work. TL, IP, NB, SB, AD, JHC, LNC, ZHG, VJ, RK, SL, KO, HO, LJO, IP, and GA initiated and designed the study, and provided funding.

## Additional information

### Accession codes

Whole genome sequence data have been deposited at the European Genome-phenome Archive (EGA, http://www.ebi.ac.uk/ega/), which is hosted by the EBI, under accession code EGAS00001000664. Genotype data for target samples is available at The database of Genotypes and Phenotypes (dbGaP, https://www.ncbi.nlm.nih.gov/gap), under accession number phs000448.v1.p1.

